# Targeting the Atypical Chemokine Receptor 2 (ACKR2) improves the benefit of anti-PD-1 immunotherapy in melanoma

**DOI:** 10.1101/2024.04.02.587761

**Authors:** Muhammad Zaeem Noman, Martyna Szpakowska, Malina Xiao, Kris Van Moer, Markus Ollert, Guy Berchem, Andy Chevigné, Bassam Janji

**Author notes:** **<u>Corresponding author:</u>** Bassam Janji (PhD, HDR), Tumor Immunotherapy and Microenvironment (TIME), Department of Cancer Research - Luxembourg Institute of Health, E mail. These authors contributed equally (Muhammad Zaeem Noman, Martyna Szpakowska, Andy Chevigné, Bassam Janji).

## Abstract

Immune checkpoint blockade (ICB) therapies, targeting PD-1 or PD-L1, have transformed cancer treatment, particularly for aggressive cancers. However, many patients fail to benefit from ICBs due to tumor characteristics, including a non-inflammatory tumor microenvironment (TME) that impedes immune cell infiltration. This study investigated the potential of targeting the Atypical Chemokine Receptor 2 (ACKR2), known for scavenging CXCR3-related chemokines crucial for lymphocyte recruitment to tumors. Genetic targeting of ACKR2 in melanoma cells increased the release of essential chemokines associated with the inflamed TME. In mouse models, ACKR2 inhibition suppressed tumor growth, improved survival, and enhanced activated immune cell infiltration into the TME. Moreover, ACKR2 targeting synergized with anti-PD-1 therapy, overcoming resistance to anti-PD-1 and improving its efficacy. Analysis of melanoma patient data from The Cancer Genome Atlas (TCGA) revealed that patients with high levels of chemokines scavenged by ACKR2 had significantly better survival rates, with increased expression of NK cell and CD8 T cell markers indicating their presence in the TME. Notably, even in patients with high CD8 expression, those expressing low ACKR2 survived better than those expressing high ACKR2. This study emphasizes the clinical importance of targeting ACKR2 as an attractive strategy for the development of combination immunotherapies to treat cold tumors, which are clinically stratified to not be eligible for ICB-based therapy.

## Short report

Cancer immunotherapies based on immune checkpoint blockades (ICB) including antibodies blocking programmed death 1 (PD-1) or programmed death ligand 1 (PD-L1) are groundbreaking treatments for several advanced and highly aggressive cancers, including melanoma, for which conventional therapies have failed. However, clinical data have revealed that relatively few patients have significant remissions from ICBs. Most have a short-term benefit or no benefit at all. To improve the benefit of ICB, therapeutic approaches are now pushing towards combining several ICBs. However, the main drawback of such approach is the cytotoxic side effects due to disruption of the fine-tuned balance of the immune system ^1^. Therefore, new combination therapeutic strategies are urgently needed to extend the use of ICBs to a large number of cancer patients and expand survival outcomes. One of the most straightforward factors responsible for the lack of responsiveness to anti-PD-1/PD-L1 is the limited infiltration of cytotoxic immune cells into the tumor microenvironment (TME) ^2^. Such infiltration strikingly relies on the establishment of an inflamed TME. According to the inflammatory status, solid tumors can be identified as “hot”, i.e., displaying a T cell-inflamed signature of pre-existing adaptive immune response and “cold” with non-T cell-inflamed signature lacking evidence of a pre-existing adaptive immune response ^3^. Therefore, therapeutic strategies that can switch “cold” tumors “hot” have inspired significant interest to improve the benefit of ICBs.

In melanoma, the presence of lymphocytes correlated with the expression of well-defined chemokine gene signature including CCL2, CCL4, CCL5, CXCL9, CCL19, CCL21, CXCL10, CXCL11, CXCL13, and XCL2. Defects in these chemokines in the melanoma TME limits the migration of activated T cells and compromises the effectiveness of antitumor immunity ^4^.

It has been shown that the newly deorphanized chemokine receptor GPR182 contributes to immunotherapy resistance in melanoma via the scavenging of CXCR3-related chemokines, reported to be important for lymphocyte recruitment to the tumour. GPR182 is primarily upregulated in melanoma-associated lymphatic endothelial cells during tumorigenesis and its knocking out increases T cell infiltration and improves antitumor immunity ^5^. Similar to GPR182, the atypical chemokine receptor 2 (ACKR2) is involved in scavenging not only numerous CC chemokines but also CXCL10 chemokine ^6,7^. Several chemokines associated with T cell-inflamed signatures, including CCL2, CCL4 and CCL5 and CXCL10 are scavenged by ACKR2 ^8^, and thus we postulated that targeting ACKR2 could be a valuable strategy to establish an inflamed TME susceptible to drive major cytotoxic immune cells and improve the benefit of anti-PD-1 immunotherapy.

To address this issue, we genetically targeted ACKR2 in B16-F10 mouse melanoma cells and assessed the release of CCL5 and CXCL10 (as a representative chemokines scavenged by ACKR2 ^6^). We showed a dramatic increase in the release of CCL5 and CXCL10 by ACKR2-targeted cells versus controls **(Figure 1A)**. We next assessed the impact of targeting ACKR2 in vivo through subcutaneous transplantation of ACKR2-defective B16-F10 cells **(Figure 1B)** in both immune-deficient NOD SCID gamma (NSG) mice lacking T, B, and NK cells as well as immuno-competent C57BL/6 mice. Our results **(Figure 1C)** showed that targeting ACKR2 had no impact on B16-F10 tumor growth and weight in NSG mice. However, a significant inhibition of tumor growth and weight was observed in ACKR2-targeted tumors in immune-competent mice, which is translated into a significant improvement of mice survival **(Figure 1D)**.

**Figure 1.**
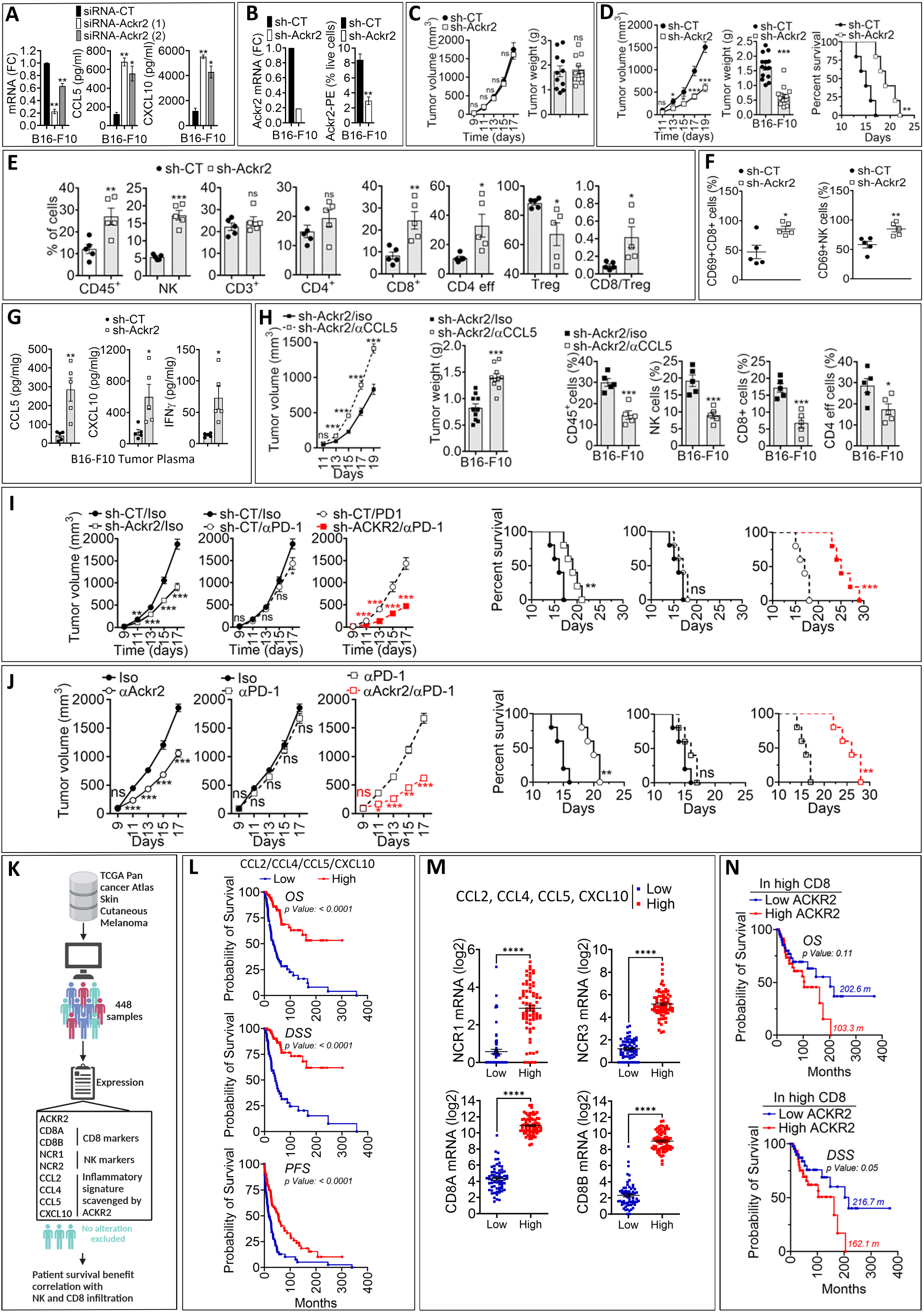
**A:** The mRNA expression of ACKR2 [reported as fold change (FC) compared to control (left)] and the release of CCL5 and CXCL10 assessed by ELISA (reported as pg/ml (right)) in mouse melanoma B16-F10 cells transfected with control siRNA (siRNA-CT) and two different ACKR2 siRNAs [siRNA-ACKR2 (1) and (2)]. **B:** The mRNA expression, reported as fold change (FC) compared to control (left), and the cell surface expression, reported as percent of live cells (right), of ACKR2 in B16-F10 cells transfected with control shRNA (sh-CT) and ACKR2 shRNAs (sh-ACKR2). **C:** Tumor growth curves (left) and weight (g) at day 17 (right) of sh-CT and sh-ACKR2 B16-F10 melanoma cells transplanted in NSG mice. Results are reported as the average of 11 mice per group as mean ± SEM (error bars). **D:** Tumor growth curves (left), tumor weight (g) at day 19 (middle), and mice survival (right) of sh-CT and sh-ACKR2 B16-F10 melanoma cells in C57BL/6 immuno-competent mice. Results are reported as the average of 15 mice per group as mean ± SEM (error bars). Mice survival curves (5 mice per group) were generated from tumor-bearing mice. A lack of survival was defined as death or tumor size >1000 mm^3^. Mice survival percentage was defined using Graph Pad Prism and P-values were calculated using the log-rank (Mantel-Cox) test (** = p≤0.01). **E:** Flow cytometry quantification of CD45+ leukocytes (gated in live cells), NK cells (NK), CD3+, CD4+, CD8+, and CD4+ effector T cells (CD4 eff) infiltrating sh-CT and sh-ACKR2 B16-F10 tumors. The results are reported as percentage (%) of cells. The last panel represents CD8+ to Treg cells ratio (CD8/Treg). **F:** Quantification of the percent of CD69+ activated CD8+ T cells and NK cells infiltrating tumors described in E. **G:** ELISA quantification of the CCL5, CXCL10 and IFNγ released in the microenvironment of sh-CT and sh-ACKR2 B16-F10 tumors. Data are reported in pg/ml standardized to excised tumor weight (g) and represented as an average of five tumors per group (each dot represents one tumor). **H:** Growth curves (left), weight in grams “g” (middle), and flow cytometry quantification of CD45+, NK, CD8+, and CD4 eff cells infiltration (right) of sh-ACKR2 B16-F10 tumors treated with control isotype (Iso) or CCL5-blocking antibody (αCCL5). I: Growth curves (left) and mice survival (right) of sh-CT and sh-ACKR2 B16-F10 tumor bearing mice treated with control isotype (Iso) or anti-PD-1 antibody (αPD-1). J: Growth curves (left) and mice survival (right) of B16-F10 tumor-bearing mice treated with control isotype (Iso) or anti-ACKR2 blocking antibody (αAckr2) alone or in combination with anti-PD-1 antibody (αPD-1). For panels I and J, mice survival curves (5 mice per group) were generated from tumor-bearing mice treated as described in I and J. A lack of survival was defined as death or tumor size >1000 mm^3^. Mice survival percentage was defined using Graph Pad Prism, and P-values were calculated using the Log-rank (Mantel-Cox) test (ns = not significant; ** = p ≤ 0.01; *** = p ≤ 0.001). For panels A to J: statistically significant differences are calculated using an unpaired two-tailed Student’s t-test. Not significant (ns) = p>0.05; * = p<0.05; ** = p<0.005; and *** = p<0.0005. K: Workflow defined for analyzing melanoma patient data described in TCGA database. **L:** Kaplan-Meier overall survival (OS, upper panel), disease-specific survival (DSS, middle panel), and progression free survival (PFS, lower panel) curves of melanoma patients expressing high and low mRNA levels of CCL2, CCL4, CCL5 and CXCL10. Patients displaying high expression of CCL2, CCL4, and CCL5 have significantly improved OS, DSS and PFS compared to those expressing low CCL2, CCL4, CCL5 and CXCL10 levels. The p-value of each curve was determined using the log-rank (Mantel-Cox) test. **M:** The mRNA expression of NK markers (NCR1 and NCR3, upper panels) and CD8 markers (CD8A and CD8B, lower panels) in melanoma patients displaying low and high expression levels of CCL2, CCL4, CCL5 and CXCL10. The results are shown as the mean ± SEM (error bars). **N:** Kaplan-Meier disease-specific survival (OS, upper panel), and overall survival (DSS, lower panels) curves of CD8 high melanoma patients expressing low and high mRNA levels of ACKR2. Probability of survival in months (m) is reported on the graphs for each group. The p-value of each curve was determined using the log-rank (Mantel-Cox) test.

Inhibition of tumor growth in ACKR2-targeted tumors was related to a significant increase in the percentage of live CD45+ cells, CD4+ effector cells, NK cells, and CD8+ T-cells associated with a decrease in Tregs **(Figure 1E)**. We also observed increased expression of the activation marker (CD69) on both NK and CD8+ T cells infiltrating ACKR2-targeted tumors **(Figure 1F)**. These results suggest that targeting ACKR2 in melanoma increased the infiltration of cytotoxic immune cells presumably by preventing the scavenging of key chemokines involved in the infiltration of immune cells such as CCL5 ^9,10^ and IFNγ. This assumption is supported by our data showing that i) higher levels of CCL5, CXCL10 and IFNγ were detected in the TME of ACKR2-targeted tumors compared to controls **(Figure 1G)**, and ii) treating mice bearing ACKR2-targeted tumors with anti-CCL5-blocking antibodies abolishes the effect of targeting ACKR2 on tumor growth inhibition and improvement of the infiltration of CD45+, NK, CD8+, and CD4+ effector cells into the TME **(Figure 1H)**.

We next evaluated the impact of targeting ACKR2 on the therapeutic benefit of anti-PD-1 using the B16-F10 melanoma model reported to be resistant to anti-PD-1. As expected, our results **(Figure 1I)** showed that anti-PD-1 monotherapy had no, or marginal, effect on B16-F10 tumor growth and mice survival. Interestingly, the therapeutic benefit of anti-PD-1 was significantly improved in mice bearing ACKR2-targeted B16-F10 tumors **(Figure 1I)** or in those bearing control B16-F10 tumors but treated with anti-ACKR2 blocking antibody **(Figure 1J)**. Such improvements were translated into a significant decrease in the tumor growth and prolonged survival **(Figure 1I and J)**. Finally, we assessed the therapeutic value of targeting ACKR2 in melanoma patients from TCGA **(Figure 1K)** by evaluating the survival benefit of those expressing high and low levels of CCL2, CCL4, CCL5 and CXCL10 chemokines, which are scavenged by ACKR2 and reported to be associated with CD8 T-cell recruitment in melanoma. Remarkably, melanoma patients expressing high levels of CCL2, CCL4, CCL5 and CXCL10 survive significantly better than those expressing low levels of these chemokines **(Figure 1L)**. Importantly, high expression levels of NK cell markers (NCR1 and NCR3) and CD8 T cell markers (CD8A and CD8B) were detected in patients with high CCL2, CCL4, CCL5 and CXCL10; this highlights their correlation with the presence of NK and CD8 T cells in the TME **(Figure 1M)**. The tangible clinical value of targeting ACKR2 is underscored by our data showing that the low expression level of ACKR2 is associated with an improved survival even in melanoma patients expressing high levels of CD8 **(Figure 1N)**.

Here, we provide experimental evidence highlighting ACKR2 as a prioritized target to switch “cold” tumors into “hot” inflamed tumors. We strongly believe that developing small molecules or neutralizing monoclonal antibodies targeting ACKR2 represents an attractive strategy for the development of combination immunotherapies to treat cold tumors, which are clinically stratified to not be eligible for ICB-based therapy. Nevertheless, further studies are needed to determine the optimal framework for the rational combination of ACKR2 inhibitors with ICB because successful combination immunotherapy approaches must be designed by considering the expression of different immune checkpoints and the infiltration of the various subset of the immune cells in the tumor microenvironment.

## Data availability

The datasets used to support the findings of this study are available from the corresponding authors upon reasonable request.

## Acknowledgments

This work was supported by grants from Luxembourg Institute of Health; Luxembourg National Research Fund (PF19/14260467/INTERceptor, BRIDGES2020/BM/15412275/SMART COMBO, BRIDGES2021/BM/16358198/TRICK-ALDH and INTER/FNRS/20/15084569/CXCL12, CORE IMPACTT C23/BM/18068832), FNRS-Televie (7.4593.19 INTERceptor, 7.4560.21 INCITE21, 7.4579.20 CD73 and 7.4559.2 IMPACT21), Roche Pharma, Fondation Recherche Cancer et Sang Luxembourg (INCOM BIOM); Action LIONS Vaincre le Cancer (AB-2020 and MZN-2020); and Stiftelsen Cancera Sweden (2022). AC and MS are part of the Marie Skłodowska-Curie Innovative Training Network ONCORNET2.0 “ONCOgenic Receptor Network of Excellence and Training” (MSCA-ITN-2020-ETN) and the European Cooperation in Science and Technology (COST) Action CA18133 European Research Network on Signal Transduction (ERNEST).

