## Supplementary information for "Targeting the Atypical Chemokine Receptor 2 (ACKR2) improves the benefit of anti-PD-1 immunotherapy in melanoma"

Noman, Szpakowska *et al.*

This file includes:

- Material and methods
- Supplementary references

**Cells and reagents**

B16-F10 cells was purchased from ATCC and cultured as described in the data sheet of ATCC and a previous report [1]. Cells were cultured in an incubator at 37°C with 5% $CO_2$ and frequently checked for mycoplasma free using Mycoalert kit (Lonza). Control and ACKR2 siRNA were purchased from ThermoFisher. Control and ACKR2 shRNA were purchased from Santa Cruz Biotechnology. Mouse CCL5/RANTES and IFN gamma DuoSet ELISA kits were purchased from R&D. $RT^2$ qPCR Primer Assay for mouse ACKR2 was purchased from Qiagen. InVivoMab anti-mouse PD-1 (CD279) (BE0273) and InVivoMab rat IgG2a isotype control (BE0089) were purchased from BioXCell (Lebanon, USA). PE-conjugated anti-ACKR2 was purchased from LSBio. Goat polyclonal anti-chemokine receptor D6 (ab1656) and Goat polyclonal IgG isotope control were purchased from Abcam. Rabbit Anti-Murine RANTES antibody (500-P118) and normal rabbit Immunoglobulin isotype were purchased from PeproTech.

**RNA extraction and SYBR Green real-time (RT)-qPCR**

Total RNA was extracted using TRIzol solution (Invitrogen) according to manufacturer's instructions. 1 µg of total RNA was treated with DNase I and converted into cDNA using TaqMan Reverse Transcription Reagent (Applied Biosystems). The mRNA expression levels were quantified by the SYBR-GREEN qPCR method (Applied Biosystems). Relative expression was calculated using a comparative Ct method (2-ΔCt). The primer sequences are available upon request.

**Tumor immune phenotyping and Flow cytometry analysis**

Tumors were harvested, mechanically dissociated into fragments (<4 mm), and enzymatically digested using a mouse tumor dissociation kit (Miltenyi Biotec) for 45 min at 37°C. Single-cell suspensions were prepared, and red blood cells were lysed by ACK (10-548E, Lonza). Cells were then counted using a Countess Automated Cell Counter (Invitrogen) and blocked for 30 minutes on ice with Fc block (TruStain fcX™ (anti-mouse CD16/32) Antibody 101320 Biolegend). Samples were stained for surface markers for lymphoid immune populations, followed by intracellular staining for FoxP3 using True-Nuclear™ Transcription Factor Buffer Set 424401 (Biolegend) according to the manufacturer's recommended protocol. The following antibodies were purchased from Biolegend: FITC anti-mouse CD45, Brilliant Violet 785 anti-mouse CD3, APC anti-mouse CD8a, APC/Fire 750 anti-mouse CD4, PE/Cy7 anti-mouse NK-1.1 antibody, Brilliant Violet 605 anti-mouse CD69, PE/Cy5 anti-mouse CD25, Brilliant Violet 421 anti-mouse FOXP3. A LIVE/DEAD Fixable Blue Dead Cell Stain Kit

(ThermoFisher Scientific) was used for viability dying. For compensation controls, single dye stains were performed and the fluorescence spread was checked using FMO controls. The level of non-specific binding was evaluated using isotype controls.

**In vivo study approval**

Animal experiments were conducted according to the European Union guidelines. The in vivo experimentation protocols were approved by the LIH ethical committee, Animal Welfare Society, and Luxembourg Ministry of Agriculture, Viticulture and Rural Development (agreements n. LECR-2018-12).

**In vivo tumor growth and mouse treatments**

C57BL/6 mice (7 weeks old) were purchased from Janvier and housed in pathogen-free conditions for one week before experiments. The mice were injected subcutaneously in the right flank with B16-F10 cells (0.2 x $10^6$ cells) diluted in 100 µl of PBS. Anti-mouse PD-1 (CD279) (BE0273) and rat IgG2a isotype control (BE0089) were diluted in InVivoPure pH 7.0 Dilution Buffer (IP0070), and administered as described in [1]. Treatment with anti-ACKR2 antibody, anti-CCL5 antibody or Rabbit Immunoglobulin isotype was performed by IP route and started at day 9 (palpable tumors) using 2.5 mg/kg in 100ul PBS (for anti-CCL5) or 75 ug per mice of anti-ACKR2 every other day until day 19. Tumor volume was measured using calipers every other day and estimated as follows: Volume ($cm^3$) = (width) 2 x length x 0.5. Mice were excluded if they did not develop tumors or developed tumors larger than the threshold defined in the approved experimentation protocols (volume > 2000 $mm^3$), as previously reported [1].

**Melanoma patient data mining**

Data from the TCGA skin cutaneous melanoma (SKCM) cohort (448 patients) were downloaded from cBioPortal (http://www.cbioportal.org/). IDs of patients displaying high and low CCL2, CCL4, CCL5 mRNA expression (z-score relative to all samples) were extracted. Each patient's vital status and survival values (overall survival and disease-specific survival) were downloaded from the TCGA database. In patients displaying high and low CCL2, CCL4, CCL5, the log2 mRNA expression level (batch normalized from Illumina HiSeq_RNASeqV2) of NK markers (NCR1 and NCR3), CD8 markers (CD8A and CD8B) and ACKR2 was identified. The differential expression of genes of interest was generated using GraphPad software. The median survival and the p-value were calculated using the log-rank (Mantel-Cox) test in GraphPad software.

### Statistical analysis

Statistical analyses were performed using GraphPad Prism 8. An unpaired two-tailed t-test was used to determine p-values between indicated groups. Results are represented as the mean ± standard error of the mean (SEM). A p-value < 0.05 was considered statistically significant ($p \leq 0.05$ = *; $p \leq 0.01$ or $\leq 0.05$ = **; $p \leq 0.001$ or $\leq 0.005$ = ***; $p > 0.05$ = not significant, ns).

### *Supplementary references*

1 Noman, M. Z. *et al.* Inhibition of Vps34 reprograms cold into hot inflamed tumors and improves anti-PD-1/PD-L1 immunotherapy. *Sci Adv* **6**, eaax7881, doi:10.1126/sciadv.aax7881 (2020).
